# Pleotropic effects of mutations in the effector domain of influenza A virus NS1 protein

**DOI:** 10.1101/390690

**Authors:** Carina F. Pereira, Helen M. Wise, Dominic Kurian, Rute M. Pinto, Maria J. Amorim, Andrew C. Gill, Paul Digard

## Abstract

**Objective:** The multifunctional NS1 protein of influenza A virus has roles in antagonising cellular innate immune responses and promoting viral gene expression. To better understand the interplay between these functions, we tested the effects of NS1 effector domain mutations known to affect homo-dimerisation or interactions with cellular PI3 kinase or Trim25 on NS1 ability to promote nuclear export of viral mRNAs.

**Results:** The NS1 dimerisation mutant W187R retained the functions of binding cellular NXF1 as well as stabilising NXF1 interaction with viral segment 7 mRNAs and promoting their nuclear export. Two PI3K-binding mutants, NS1 Y89F and Y89A still bound NXF1 but no longer promoted NXF1 interactions with segment 7 mRNA or its nuclear export. The Trim25-binding mutant NS1 E96A/E97A bound NXF1 and supported NXF1 interactions with segment 7 mRNA but no longer supported mRNA nuclear export. Analysis of WT and mutant NS1 interaction partners identified hsp70 as specifically binding to NS1 E96A/E97A. Whilst these data suggest the possibility of functional links between NS1’s effects on intracellular signalling and its role in viral mRNA nuclear export, they also indicate potential pleiotropic effects of the NS1 mutations; in the case of E96A/E97A possibly via disrupted protein folding leading to chaperone recruitment.

## Introduction

Influenza A virus (IAV) infects both man and animals, including species important for food security [1]. A better understanding of its replication cycle is important to aid control measures. Like all viruses, IAV co-opts multiple cellular systems for replication and also has to counteract innate immune defence mechanisms. A key virally-encoded factor in both processes is the NS1 protein. As a nuclear-replicating virus, IAV utilises the NXF1 cellular mRNA export pathway to direct several of its mRNAs to the cytoplasm [2] and we and others have recently found that NS1 acts as an adaptor between segment 7 mRNAs (encoding the M1 or M2 proteins) and cellular mRNA processing/export pathways [3, 4]. NS1 is also the primary viral antagonist of innate immune responses, interfering with the function of multiple cellular polypeptides including PKR, 2’-OAS, CPSF30 and Trim25 [5]. NS1 also activates the phosphoinositide 3-kinase (PI3K) complex, possibly to inhibit apoptosis [6]. Structurally, NS1 consists of an N-terminal RNA-binding domain and a C-terminal “effector” domain followed by a disordered tail, with no identified enzymatic activities. Instead, it functions by binding to multiple different proteins and RNAs (viral and cellular) and altering their activities [5]. Various NS1 point mutations have been identified that inhibit these interactions and comparative analyses of the behaviour of wild type (WT) and mutant IAV strains with these alterations have provided important information on the biological significance of each particular function. Here, we tested whether mutations identified as affecting NS1 interactions with the p85ß subunit of PI3K (Y89A and Y89F), Trim25 (E96A/E97A) and NS1 effector domain-mediated oligomerisation (W187R) [7–11] also affected NS1’s ability to promote nuclear export of segment 7 viral mRNAs.

## Methods

All IAV plasmid constructs and viruses were based on the A/PR/8/34 (PR8) strain described previously [12]. NS1 mutations used in this study were introduced by standard PCR mutagenesis reactions; primer sequences are available on request. Methods for virus reverse genetics, infection, minireplicon assays, fluorescent *in situ* hybridization (FISH), immunofluorescence and confocal microscopy, GFP-trap pulldowns, western blots and reverse transcriptase-radioactive primer extension are described elsewhere [4]. LC-MS analysis was performed using a RSLCnano system (Thermo Fisher Scientific) coupled to a micrOTOF QII mass spectrometer (Bruker) on in-gel trypsin digested protein bands excised from Coomassie Blue-stained SDS-PAGE gels [13]. Raw spectral data were processed to peak lists and searched using Mascot 2.4 server (Matrix Science) against the Uniprot Human sequence database containing 93,786 entries.

## Results

Our previous work led to the hypothesis of NS1 acting as an adaptor protein to feed viral segment 7 mRNA(s) into the cellular NXF1 export pathway, by forming a ternary complex between NS1, NXF1 and the mRNA that requires both the NS1 effector domain and a functional RNA-binding domain [4]. To further test the role of the NS1 effector domain, a set of point mutations in this region of the protein were tested (Table 1), firstly in a subviral “minireplicon” system. 293T cells were transfected with plasmids to produce segment 7 ribonucleoproteins (RNPs) as well as with either WT or mutant NS1s as EGFP-fusion proteins. Cells were fixed 24 h post transfection and segment 7 mRNA localisation detected by FISH (**Figure 1A**). NS1 expression was confirmed by detection of GFP signal as well as by western blot, although the E96A/E97A mutant accumulated to lower levels than the other NS1s (**Figure 1B**). WT NS1-GFP and most mutant proteins localised largely to the cytoplasm, but the W187R mutant was found both in nucleus and cytoplasm (**Figure 1A**). No mRNA staining was detected in the negative control lacking a complete RNP. As expected [4], when the minimal components for RNP reconstitution were present, segment 7 mRNA was detected primarily in cell nuclei, whilst on addition of NS1-GFP, it translocated to the cytoplasm. However, in the presence of PI3K-binding mutants NS1-Y89A or NS1-Y89F or the Trim25-binding mutant NS1-E96A/E97A, segment 7 mRNA was clearly still retained in the nucleus. In contrast, the dimerization mutant NS1-W187R caused segment 7 mRNA to localise to the cytoplasm. To quantify these effects, cells were visually scored according to the predominant cellular localisation of segment 7 mRNA (nuclear, cytoplasmic or both). In the absence of NS1, segment 7 was mainly nuclear in ~80% of cells scored, while the remaining cells presented mRNA staining both in the nucleus and the cytoplasm (**Figure 1C**). When WT NS1 or NS1-W187R were added, a statistically significant shift of segment 7 mRNA localisation was confirmed, with ~ 75% of cells showing mRNA in the cytoplasm. However, addition of NS1-E96A/E97A, NS1-Y89A or NS1-Y89F did not produce this shift, with a clear majority of cells displaying nuclear segment 7 mRNA, similarly to samples with no NS1-GFP.

**Table 1.** Genetic and phenotypic summary of the NS1 polypeptides used in this study.

| NS1 protein | mRNA export | NXF1 binding | M1 mRNA binding | Functional alteration | Reference |
| --- | --- | --- | --- | --- | --- |
| WT | + | + | + | n/a | [12] |
| N81<sup>a,b</sup> | - | - | - | Effector domain deletion | [14] |
| R38A/K41A<sup>b</sup> | - | - | - | dsRNA-binding | [15] |
| F103S/M106I<sup>b</sup> | + | + | + | Gain of CPSF inhibition | [16] |
| Y89A | - | + | - | PI3K-binding | [17] |
| Y89F | - | + | - | PI3K-binding | [8] |
| E96A/97A | - | + | + | Trim25-binding | [9] |
| W187R | + | + | + | Homo-dimerisation | [11] |

<sup>a</sup>Stop codon added at position 82. <sup>b</sup>See [4] for experimental data.

**Figure 1.**
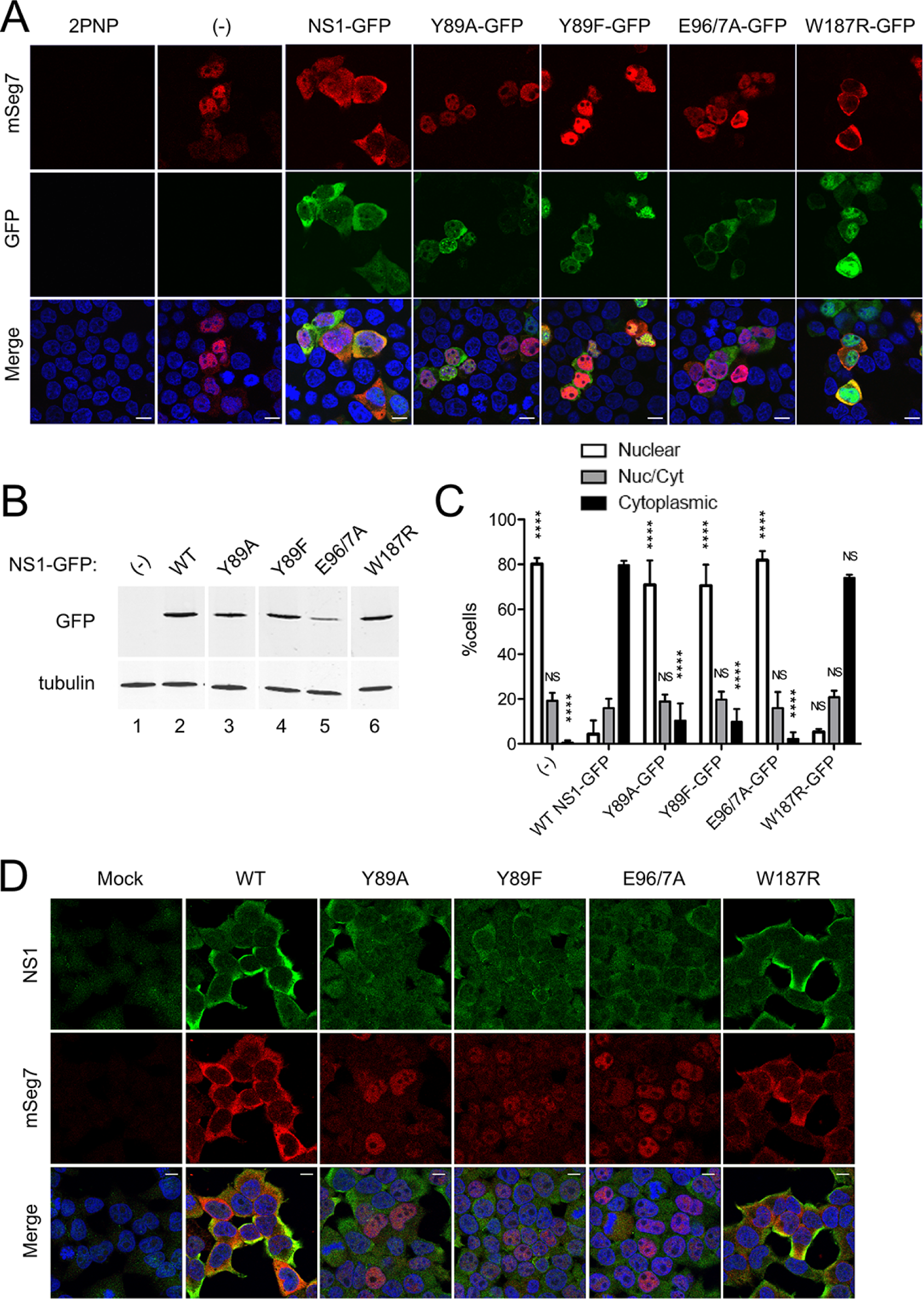
Effect of mutations in the NS1 effector domain on segment 7 mRNA localisation. A-C) 293T cells were transfected with plasmids to recreate segment 7 RNPs or, as a negative control with a combination that excluded a plasmid expressing PB2 (2PNP), together with the indicated NS1-GFP polypeptides. (A) Cells were processed for FISH analysis at 24 h post transfection (p.t.) using Cy3-labelled RNA probes for detection of positive sense segment 7 viral RNAs (red). GFP fluorescence was also detected (green). Images were captured using a Leica-TCS confocal microscope and Leica TCS analysis software. (B) Duplicate cell samples were lysed and examined by SDS-PAGE and western blotting for GFP and tubulin. (C) Individual cells were scored according to the predominant cellular localisation of segment 7 mRNA considering three phenotypes: nuclear, cytoplasmic or mixed. Values are the mean ± range from two independent experiments. A 2-way ANOVA was used to test for statistically significant differences between the values for each category from those obtained for WT-NS1 (NS, not significant; **** P < 0.0001). (D) 293T cells were mock infected or infected with the indicated viruses at an M.O.I. of 5. At 6 h post infection (p.i.), cells were processed for FISH as above (red) and stained with anti-NS1 serum (green) and DAPI (for DNA; blue). Scale-bars indicate 10 µm. Lanes 1 and 2 in panel (B) and a portion of the corresponding numerical data in panel (C) are taken with permission from **Figure 4C** and 4B respectively of [4].

To further test the mRNA export activity of the NS1 effector domain mutants, their phenotype was examined in the context of virus infection. 293T cells were mock infected or infected with either WT PR8 or the NS1 mutant viruses and at 6 h post infection (p.i.), fixed and processed for segment 7 FISH analysis as well as immunofluorescently stained for NS1 (**Figure 1D**). All viruses generally gave cytoplasmic NS1 signal above the uninfected cell background, although staining of cells infected with NS1-Y89 and NS1-E96A/E97A mutants was noticeably fainter than the WT and W187R viruses. No viral mRNA signal was detected in the mock sample while segment 7 mRNA was cytoplasmic in WT and W187R virus-infected cells. In cells infected with NS1-Y89A, NS1-Y89F and NS1-E96A/E97A, segment 7 mRNA was predominantly nuclear. Thus, in both minireplicon and viral settings, the NS1 dimerization mutant W187R supported normal segment 7 mRNA nuclear export, but the PI3K-binding and Trim25-binding mutants showed loss-of-function.

NS1 both binds NXF1 and promotes the ability of NXF1 to bind segment 7 mRNA [4, 18–20]. To test whether the effector domain mutants retained NXF1-binding activity, 293T cells were transfected with either GFP or GFP-NXF1 and 48 h later mock infected or infected with the panel of viruses. At 6 h p.i., cells were lysed and GFP-Trap precipitations were performed on the supernatants. Western blot analyses of total and bound fractions showed that GFP and GFP-NXF1 were both expressed and collected in comparable amounts (**Figure 2A**). Blotting for NS1 confirmed successful infection, although the NS1-E96A/E97A polypeptide again accumulated to lower levels than the other NS1s. Nevertheless, all NS1 polypeptides co-precipitated with GFP-NXF, indicating that the mutations did not abrogate the interaction.

**Figure 2.**
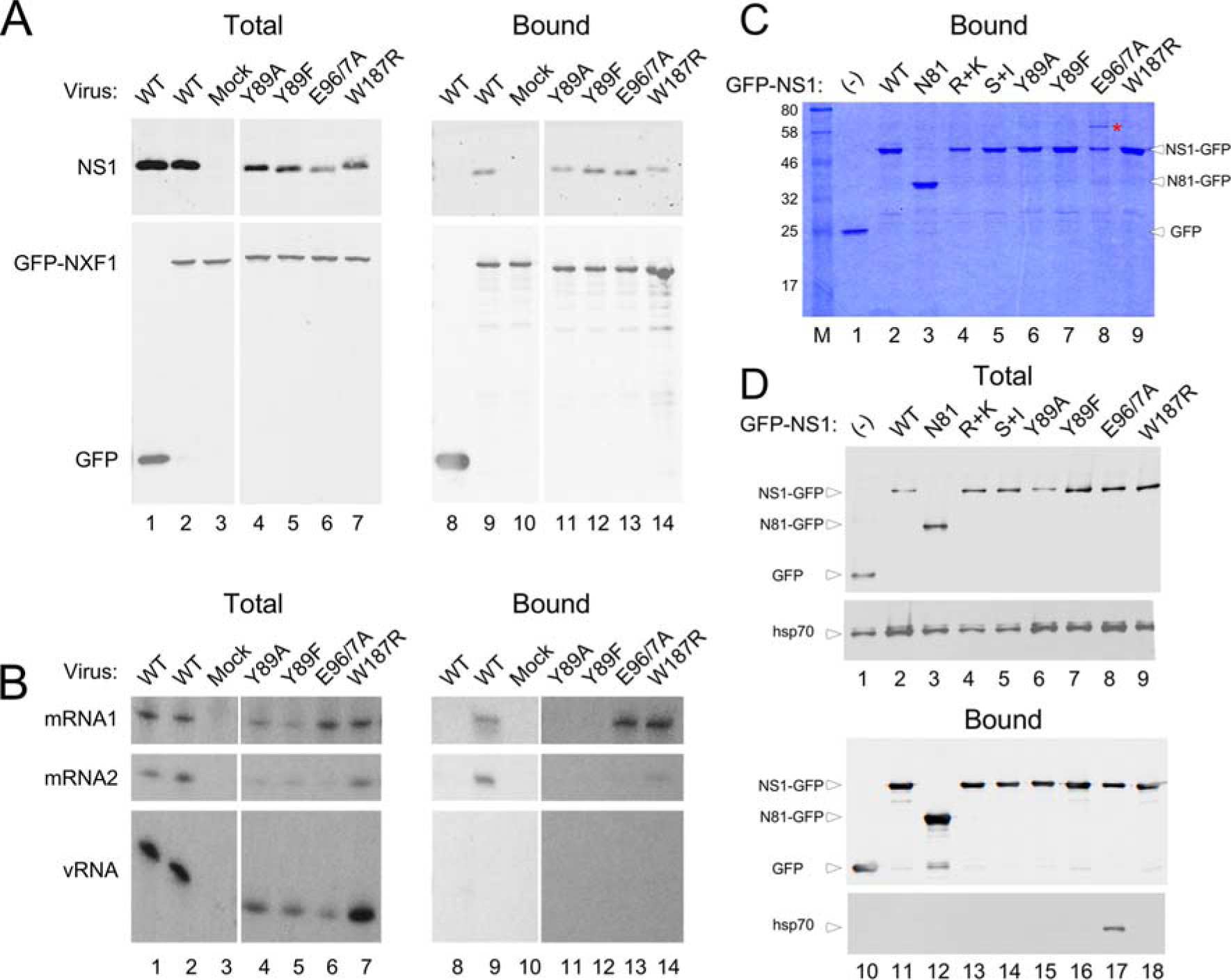
Interactions between NS1, cellular proteins and segment 7 mRNAs. (A) 293T cells were transfected with plasmids encoding GFP or GFP-NXF1 and 48 h later either mock infected or infected with the indicated NS1 mutant viruses at an M.O.I. of 5. Cells were harvested at 6 h p.i. and cell lysates examined by western blotting for the indicated proteins before (Total) or after (Bound) being subjected to GFP-Trap pulldown assays. (B) 293T cells were transfected with GFP-NXF1 (GFP-NXF1 +) or with GFP alone (GFP-NXF1 -) and 48 h later either mock infected or infected with the indicated viruses at an M.O.I. of 10, in duplicate sets. At 6 h p.i., cells were harvested and total cellular RNA was extracted from one set while the second set was subjected to GFP-Trap pulldown assays prior to total RNA extraction. RNA was analysed by reverse transcription with radiolabelled primers specific for segment 7 RNAs followed by urea-PAGE and autoradiography. (C, D) 293T cells were transfected with either GFP alone or with GFP-tagged NS1 mutant protein expressing plasmids as indicated and 48 h later cells were harvested and subjected to GFP-Trap pull down assays. Eluted samples were loaded onto SDS-PAGE gels and (C) stained with Coomassie Blue dye or (D) western blotted for GFP or hsp70 before (Totals) or after (Bound) GFP-trap fractionation. Lanes 1-3 and 8-10 from panel (A) are reproduced with permission from Figure 7 of [4].

To test whether the effector domain mutations affected the ability of NS1 to promote a stable interaction between NXF1 and segment 7 mRNA, 293T cells were transfected with GFP or GFP-NXF1 and infected with WT or mutant viruses as above. At 6 h p.i., cells were lysed and total RNA was extracted before or after the lysates had been subjected to GFP-Trap pulldown. Segment 7 RNA species were then detected by primer extension. Viral mRNAs and genomic vRNA were detected in the total fraction of every infected sample (**Figure 2B**, lanes 1-2 and 4-7). Analysis of the bound fraction from WT virus infected cells showed that both segment 7 mRNA species but not genomic vRNA co-precipitated with GFP-NXF1 (lane 9) while no viral RNAs bound to GFP alone (lane 8), consistent with previous work [4, 18] In contrast, viral mRNAs were not detected in GFP-NXF1 pulldowns from cells infected with either NS1-Y89A or NS1-Y89F viruses (lanes 11 and 12). The NS1-W187R and -E96A/E97A mutants also supported NXF1 interactions with the viral mRNAs (lane 13, 14). Since the E96A/E97A mutation blocked successful nuclear export of segment 7 mRNA, this finding indicated that, like the interaction between NS1 and NXF1, the interaction between NXF1 and segment 7 is necessary, but not sufficient for the successful nuclear export of segment 7 mRNA.

The behaviour of the NS1 E96A/E97A mutant – unable to mediate nuclear export of segment 7 mRNA despite binding to NXF1 and promoting its interaction with the mRNA – suggested the involvement of another essential factor in the hypothesised ternary export complex. Accordingly, we examined cellular interaction partners of WT NS1 and the mutants, including a complete deletion of the effector domain (NS1-N81), an RNA-binding domain mutant (R38A/K41A) and a gain-of-function mutant that restores CPSF30-binding activity (F103S/M106I), of which only the latter supported mRNA nuclear export (Table 1; [4]). 293T cells were transfected with GFP or GFP-tagged NS1 mutant plasmids. Two days later, cells were collected, lysed and subjected to GFP-Trap. Bound fractions were loaded onto SDS-PAGE gels and stained with Coomassie Blue. No obvious contamination from cellular GFP-binding proteins was seen from cells transfected with the GFP control plasmid (**Figure 2C**, lane 1). Major polypeptide species of the expected size were detected in all other samples, including the truncated form NS1-N81, confirming successful expression (lanes 2-9). Various other minor polypeptide species were also visible that were mostly common to all NS1-containing samples. However, one clear difference was observed; a distinct novel polypeptide in the E96A/E97A preparation (lane 8) that was absent from other samples. LC-MS analysis of the excised band unambiguously identified this polypeptide as heat shock protein 70 (hsp70) through multiple peptide matches (**Table S1**).

To confirm the interaction between NS1-E96A/E97A and hsp70, repeat GFP-trap pulldowns were analysed by western blotting for hsp70 as well as for the GFP-tagged “bait” proteins. Transfection was confirmed by the detection of GFP and GFP-tagged NS1 mutant proteins, while equivalent amounts of the cellular proteins hsp70, Nup62 and UAP56 in the total fractions of all samples confirmed that comparable amounts of cell lysates had been generated (**Figure 2D**, lanes 1-9). However, in the bound fraction, hsp70 was detected only in the NS1-E96A/E97A sample (lane 17), confirming the specific interaction with hsp70 for NS1 E96A/E97A.

## Discussion

The data reported here extend our previous investigation [4] into the role of NS1 in facilitating nuclear export of viral mRNA. Firstly, we show that higher order multimerization of NS1 via its effector domain is not needed for viral mRNA nuclear export. Secondly, the finding that point mutations in the NS1 effector domain, and not just its complete deletion along with the C-terminal unstructured region [4], can block viral mRNA nuclear export, strengthens the argument for the functional involvement of this domain in the process. Thirdly, identification of an NS1 mutant (E96A/E97A) that still binds NXF1 and enhances its interaction with segment 7 mRNA without upregulating mRNA nuclear export suggests that at least one other factor is needed for export. Finally, the effect of mutations previously considered to specifically inhibit PI3K activation or inhibition of RIG-I signalling on M1 mRNA localisation could suggest the involvement of one or both of these pathways in viral mRNA nuclear export.

Of broader significance than the mechanism of IAV mRNA nuclear export, our data add to the evidence suggesting pleiotropic effects from NS1 mutations commonly used as tools to interrogate viral interactions with the PI3K and RIG-I pathways (e.g. [10, 17, 21–28]. Potential pleiotropic effects of the E96A/E97A mutation have been noted previously, including poor expression of the polypeptide as seen here [25] and an intriguing effect on PI3K activation [29]. Indeed a recent report detailing the crystal structure of NS1-Trim25 complexes concluded that NS1 residues E96 and 97 do not directly contact Trim25 but instead support a network of contacts within the local region of the effector domain in which L95 and S99 actually make contact with Trim25 [30]. Our finding that the E96A/E97A mutation also induces binding of hsp70 further suggests effects on protein folding and that interpreting results obtained with this mutation may be complicated.

## Limitations

Attempts to probe the role of PI3K activation in IAV mRNA export using small molecule inhibitors gave inconsistent results (data not shown) and could not be further pursued because of time and funding constraints. It would also have been interesting to test the effects of hsp70 inhibitors [31] had time allowed. The effect of NS1 mutations L95A/S99A on mRNA export should also be tested.

## Declarations

### Ethics approval and consent to participate

not applicable.

### Availability of data and material

available from the corresponding author on reasonable request.

## Acknowledgements

we thank Professor Adrian Whitehouse for the gift of pGFP-NXF1.

## Consent for publication

not applicable.

## Author’s contributions

CFP carried out the majority of the experimental work with input from MJA, HW and RMP. DK and ACG performed the LC-MS analyses. CFP and PD drafted the manuscript and all authors read, corrected and approved the final manuscript.

